# SequencEnG: an Interactive Knowledge Base of Sequencing Techniques

**DOI:** 10.1101/319079

**Authors:** Yi Zhang, Mohith Manjunath, Yeonsung Kim, Joerg Heintz, Jun S. Song

## Abstract

Next-generation sequencing (NGS) techniques are revolutionizing biomedical research by providing powerful methods for generating genomic and epigenomic profiles. The rapid progress is posing an acute challenge to students and researchers to stay acquainted with the numerous available methods. We have developed an interactive online educational resource called SequencEnG (acronym for <u>Sequenc</u>ing Techniques <u>En</u>gine for <u>G</u>enomics) to provide a tree-structured knowledge base of 66 different sequencing techniques and step-by-step NGS data analysis pipelines comparing popular tools. SequencEnG is designed to facilitate barrier-free learning of current NGS techniques and provides a user-friendly interface for searching through experimental and analysis methods. SequencEnG is part of the project KnowEnG (Knowledge Engine for Genomics) and is freely available at http://education.knoweng.org/sequenceng/.

## 1 Introduction

The bioinformatics field is constantly challenged with the increasing diversity and large amounts of experimental data being generated. For instance, numerous high-throughput next-generation sequencing (NGS) techniques, such as RNA-seq, ChIP-seq, and Hi-C, are now widely used in biomedical research, each with accompanying active development of computational algorithms and tools (Shendure *et al.*, 2017). Students and researchers are thus challenged with the difficulty of keeping pace with the rapidly growing number of new NGS techniques and their analysis methods. The research community as a whole would therefore benefit from a centralized resource to learn about currently available NGS techniques. There are already some resources available to explore NGS data analysis tools, such as OMICtools (Henry *et al*., 2014) and AZTEC (https://aztec.bio/). OMICtools is an online database of bioinformatics tools where a bioinformatics software developer can submit their own tool and a researcher can discover these tools. The number of tools listed in the OMICtools database, where tool evaluation is based on user reviews, has been steadily increasing. However, this resource relies on active user participation, and many listed tools currently have no reviews or ratings available. AZTEC is another web resource where many bioinformatics tools, web services, publications and libraries are documented; but, it currently does not provide self-contained educational resources or basic analysis pipelines. Other web resources contain data analysis tutorials, but are only for a specific analysis; e.g., GATK (https://software.broadinstitute.org/gatk/) for variant calling and Homer (http://homer.ucsd.edu/homer/basicTutorial/index.html) for motif, ChIP-seq and RNA-seq analyses. The above resources also lack an overview and comparison among commonly used and state-of-the-art tools.

The National Institute of Health (NIH) has recently funded Big Data to Knowledge (BD2K) Centers as well as multiple BD2K Training projects to develop and disseminate educational resources for bioinformatics. As part of the Knowledge Engine for Genomics (KnowEnG) BD2K center (Kim *et al*., 2017; Manjunath *et al.,* 2018), we have developed a web resource called SequencEnG that annotates and explains currently available NGS techniques and their data analysis methods. Our interactive website consists of two parts: the NGS knowledge tree, and the data analysis pipelines. SequencEnG contains 66 NGS techniques and organizes them into a knowledge tree based on genetic or epigenetic information being assayed. For each sequencing technique, we graphically illustrate the major experimental steps involved in the protocol. The interactive data analysis pipelines are currently available for 13 popular NGS techniques, including RNA-seq, ChIP-seq, Hi-C, and Whole Genome Bisulfite Sequencing (WGBS). For each analysis step in the pipeline, major bioinformatics tools are listed in a searchable table containing the description, comparison to similar tools, evaluation from benchmarking papers, and other relevant features.

## 2 Software description

### 2.1 A knowledge tree of sequencing techniques

We have manually curated 66 NGS techniques with distinct sequencing goals, amounting to 3 general and 27 specific categories classified as branches in an interactive knowledge tree (Fig. 1A). Each leaf of the tree shows one NGS technique with a corresponding description that references the original NGS paper and includes a simple yet informative graphical summary of the experimental steps (Fig. 1A, bottom right). An interactive tutorial will take the user on a tour of the website when the user visits the SequencEnG website for the first time.

**Fig. 1.**
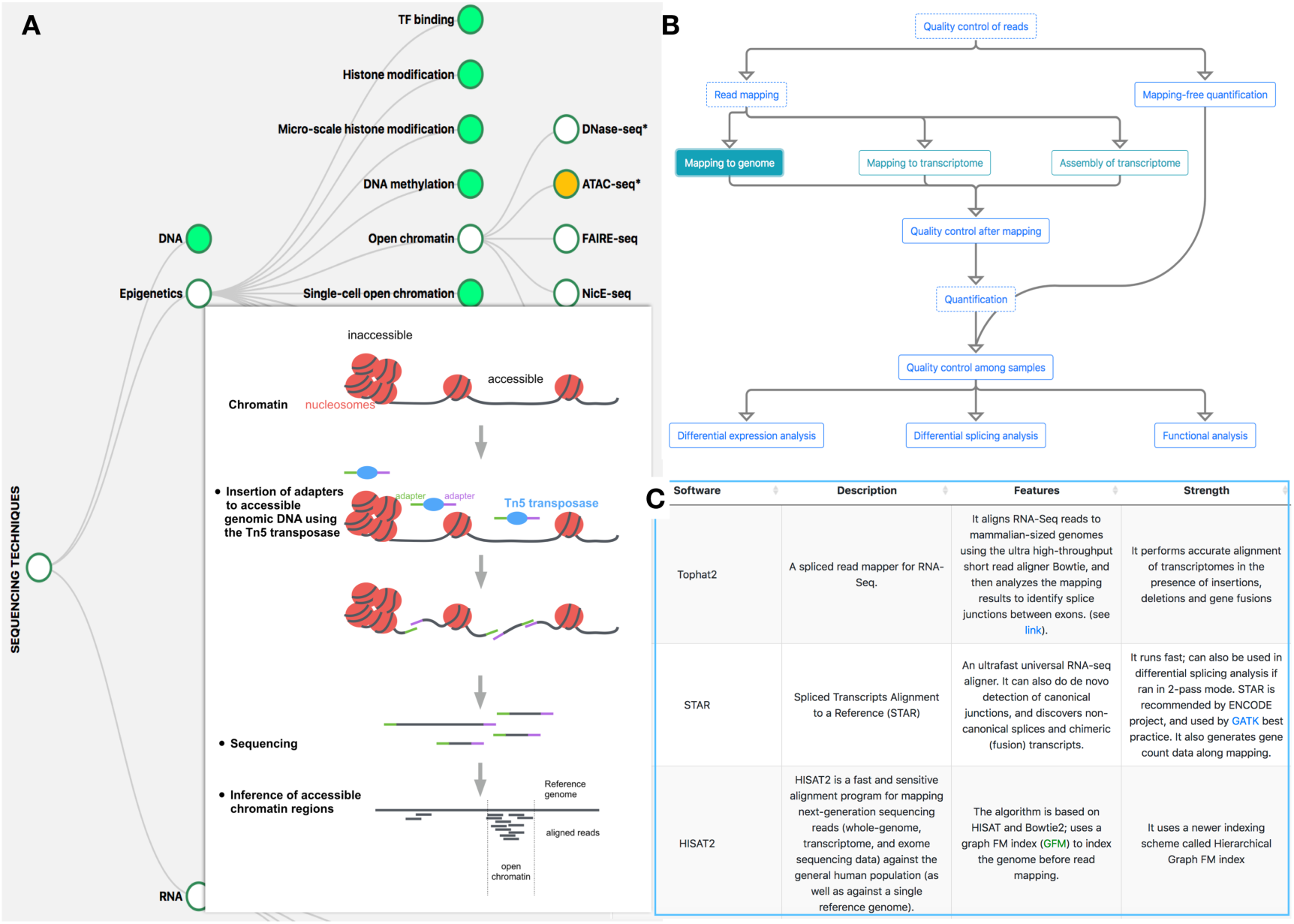
Snapshots of SequencEnG. (**A**) The interactive NGS knowledge tree with sequencing goals as branches and sequencing techniques as leaves; Bottom right shows an example of the graphical experimental steps for ATAC-seq. (**B**) An example of the analysis pipeline for RNA-seq. (**C**) An example of major bioinformatics tools available.

### 2.2 Analysis pipelines with tool comparisons

For 13 of the most widely used NGS techniques, we provide interactive analysis pipelines (Fig. 1B) and corresponding popular computational tools available for each step in the pipeline (Fig. 1C). For example, different pipelines for performing RNA-seq analysis, from quality control to differential gene expression calling, are illustrated in a flowchart in Fig. 1B, where gene quantification can be done with canonical read mapping methods such as HISAT (Kim *et al*., 2015), or with alignment-free tools such as kallisto (Bray *et al.*, 2016).

A set of popular tools for each pipeline step is shown in a searchable table when the user selects a step. The table summarizes their basic features, strengths, limitations, input/output format, working platforms, references, and other features such as whether a ChIP-seq peak caller is suitable for transcription factors or histone modifications, and whether a tool accepts specific file formats from another software. For example, sleuth (Pimentel *et al.*, 2017) accepts the output of kallisto. The respective strengths and limitations are summarized from available benchmarking papers or resources, with references and links provided in the interactive table. The table thus not only includes multiple tools available, but also provides evaluation information in three ways: 1) performance evaluation based on benchmarking and review papers; 2) application in large projects such as ENCODE (ENCODE Project Consortium, 2012); 3) ranking based on Relative Citation Ratio (RCR, Hutchins *et al.*, 2016) score queried from iCite (https://icite.od.nih.gov/) to reflect the importance and popularity of the tool. Tool recommendation and comparison can be a major challenge in the rapidly growing field of bioinformatics for various reasons; for example, it is difficult to assess the quality of algorithms for calling topologically associated domains (TADs) in Hi-C data analysis, because of the ambiguity in defining the notion of TAD in biology (Forcato *et al.*, 2017). Therefore, instead of recommending a single tool per pipeline step, SequencEnG allows researchers to be aware of various strengths and limitations of using popular tools.

## 3 Conclusions

SequencEnG is an interactive web resource for active learning or teaching NGS techniques, graphically summarizing experimental protocols and describing detailed analysis pipelines. The user can interact with the NGS knowledge tree to learn about 66 different NGS techniques, and also explore the NGS data analysis pipelines to gather quickly a list of popular and useful tools. SequencEnG thus provides a friendly learning environment for students and researchers entering bioinformatics and the NGS field, especially for those in small colleges without extensive teaching resources on these subjects. While continuing our manual curation, we will explore automating future updates by utilizing the strength of text mining in our BD2K Center. To help the user be aware of new developments in the sequencing field, the Resources page of SequencEnG contains useful links and interactive pipelines from ENCODE, Cistrome (Liu *et al.*, 2011), and Genomic Data Commons (Grossman *et al.*, 2016).

## Acknowledgements

The authors would like to thank the members of the Song group for feedback.

## References

Shendure, J., Balasubramanian, S., Church, G. M., Gilbert, W., Rogers, J., Schloss, J. A., & Waterston, R. H. (2017). DNA sequencing at 40: past, present and future. Nature, 550(7676), 345.

Henry, V. J., Bandrowski, A. E., Pepin, A. S., Gonzalez, B. J., & Desfeux, A. (2014). OMICtools: an informative directory for multi-omic data analysis. Database, 2014.

Kim, M., Kim, Y., Qian, L., & Song, J. S. (2017) TeachEnG: a Teaching Engine for Genomics. Bioinformatics, 33, 3296–3298.

Manjunath, M., Zhang, Y., Kim, Y., Yeo, S. H., Sobh, O., Russell, N., Followell, C., Bushell, C., Ravaioli, U. & Song, J. S. (2018). ClusterEnG: an interactive educational web resource for clustering and visualizing high-dimensional data. PeerJ Computer Science, 4, e155.

Kim, D., Langmead, B., & Salzberg, S. L. (2015). HISAT: a fast spliced aligner with low memory requirements. Nature methods, 12(4), 357.

Bray, N. L., Pimentel, H., Melsted, P., & Pachter, L. (2016). Near-optimal probabilistic RNA-seq quantification. Nature biotechnology, 34(5), 525.

Pimentel, H., Bray, N. L., Puente, S., Melsted, P., & Pachter, L. (2017). Differential analysis of RNA-seq incorporating quantification uncertainty. Nature methods, 14(7), 687.

ENCODE Project Consortium. (2012). An integrated encyclopedia of DNA elements in the human genome. Nature, 489(7414), 57.

Hutchins, B. I., Yuan, X., Anderson, J. M., & Santangelo, G. M. (2016). Relative Citation Ratio (RCR): A new metric that uses citation rates to measure influence at the article level. PLoS biology, 14(9), e1002541.

Forcato, M., Nicoletti, C., Pal, K., Livi, C. M., Ferrari, F., & Bicciato, S. (2017). Comparison of computational methods for Hi-C data analysis. Nature methods, 14(7), 679.

Liu, T., Ortiz, J.A., Taing, L., Meyer, C.A., Lee, B., Zhang, Y., Shin, H., Wong, S.S., Ma, J., Lei, Y. & Pape, U.J. (2011). Cistrome: an integrative platform for transcriptional regulation studies. Genome biology, 12(8), p.R83.

Grossman, R.L., Heath, A.P., Ferretti, V., Varmus, H.E., Lowy, D.R., Kibbe, W.A., & Staudt, L.M. (2016). Toward a shared vision for cancer genomic data. New England Journal of Medicine, 375(12), pp.1109–1112.

